# Two different phospholipases C, Isc1 and Pgc1, cooperate to regulate mitochondrial function

**DOI:** 10.1101/2022.03.18.484854

**Authors:** Balazova Maria, Babelova Lenka, Durisova Ivana, Vesela Petra, Kanovicova Paulina, Zahumensky Jakub, Malinsky Jan

## Abstract

Absence of Isc1, the yeast homologue of mammalian neutral sphingomyelinase type 2, leads to severe mitochondrial dysfunction. We show that deletion of another type-C phospholipase, the phosphatidylglycerol (PG)-specific Pgc1, rescues this defect. Phosphatidylethanolamine (PE) levels and cytochrome *c* oxidase activity, reduced in *isc1*Δ cells, were restored to wild-type levels in the *pgc1*Δ*isc1*Δ mutant. Pgc1 substrate, PG, inhibited *in vitro* activity of Isc1 and phosphatidylserine decarboxylase Psd1, an enzyme crucial for PE biosynthesis. We also identify a mechanism by which the balance between the current demand for PG and its consumption is controlled. We document that the product of PG hydrolysis, diacylglycerol, competes with the substrate of PG-phosphate synthase, Pgs1, and thereby inhibits the biosynthesis of excess PG. This feedback loop does not work in the absence of Pgc1, which catalyzes PG degradation. Finally, Pgc1 activity is partially inhibited by products of Isc1-mediated hydrolysis. The described functional interconnection of the two phospholipases contributes significantly to lipid homeostasis throughout the cellular architecture.

**Summary:** The coordinated action of two different type-C phospholipases is documented, which provides a balance between mitochondrial phospholipid biosynthesis and sphingolipid metabolism. The regulatory role of specific lipids, phosphatidylglycerol, diacylglycerol and ceramide in this process is demonstrated.

## Introduction

Together with glycerophospholipids and sterols, sphingolipids represent one of the main classes of lipids forming the biological membranes of all eukaryotic organisms. Besides their essential structural function, sphingolipids and their precursors, long chain bases and ceramides, participate in cell signaling during polarized growth, cell cycle control, adaptation to various types of environmental and ER stresses, etc. (1 and references therein; 2–3). In higher eukaryotes including humans, dysregulated sphingolipids and especially ceramides underlie severe pathologies including type-2 diabetes, cardiovascular diseases or cancer (e.g. 4–6). In cellular metabolism, ceramides are generated either during sphingolipid neosynthesis or by the hydrolysis of complex sphingolipids.

In yeast, complex sphingolipids (inositolphosphoceramides and their mannosylated variants) are hydrolyzed to ceramides by a single inositolphosphosphingolipid phospholipase C, Isc1, an enzyme homologous to mammalian neutral sphingomyelinase type 2 (7, 8). During fermentation, Isc1 localizes to the membrane of endoplasmic reticulum. During the diauxic shift the protein changes its localization towards the outer mitochondrial membrane (9). Translocation of Isc1 to mitochondria and increase of its activity is induced following the protein phosphorylation by Sch9 kinase (10).

Sch9 itself requires dual phosphorylation to activate its kinase function, combining signals from the membrane stress-detecting eisosome complex at the plasma membrane (11, 12) and nutrient-responding TORC1 at the membrane of vacuole (13, 14). Eisosomal branch of this signaling pathway includes Pkh1/2 kinases and the eisosome core protein Pil1 (15, 16). Sch9 activity stabilizes the ceramide pool through a combined action at sphingolipid degradation and neosynthesis pathways. Specifically, it not only stimulates Isc1-mediated hydrolysis of complex sphingolipids, but also represses expression of genes coding ceramidases *YDC1* and *YPC1* (10). The importance of this balanced regulation is well illustrated by the data obtained in *isc1Δ* cells, in which both TORC1- and Pkh1-mediated phosphorylations of Sch9 contribute to their respiration defects (17, 18; respectively).

Ceramides generated by Isc1 contribute to normal mitochondrial function. The increased Isc1 activity in cells after the diauxic shift positively correlates with increased levels of the product of Isc1-catalyzed complex sphingolipids hydrolysis, phytoceramide, in these cells. At the same time, both effects are conditioned by running phosphatidylglycerol (PG) synthesis in mitochondria (19). PG is a tightly regulated, low-abundant phospholipid in yeast, predominantly serving as a precursor in cardiolipin (CL) biosynthetic pathway. Pgs1-catalyzed synthesis of PG-phosphate (PGP), which takes place at the inner mitochondrial membrane, represents a rate-limiting step of the pathway. Following the biosynthesis, PG is either utilized as a substrate by CL synthase Crd1, or spread to other cellular membranes, where it is subject to rapid degradation by a specific phospholipase C, Pgc1 (20).

Supply of exogenous phytoceramide was enough to rescue the growth defect of the cells lacking either Isc1 or PGP synthase, Pgs1 (19). However, at least in the latter strain, this could not be related to rescue of the mitochondrial function because PG and CL, anionic phospholipids indispensable for respiration (21), were still absent. Anyway, the excess ceramide was somehow able to participate in improving the phenotype of not only *isc1*Δ, but also *pgs1*Δ cells. As mentioned above, in wild-type diauxic cells, the increased ceramide content depends on PG synthesis (19). In this study, we asked whether elevated PG levels themselves could at least partially rescue the phenotype of *isc1*Δ cells. Deletion of *PGC1* gene has been shown before to induce PG accumulation in various backgrounds (20, 22, 23). Therefore, we constructed *pgc1*Δ*isc1*Δ double deletion mutant. Detailed characterization of this double deletion strain reveals a tight interconnection between mechanisms of sphingolipid and phospholipid metabolism regulation.

## Results and Discussion

### *Respiration defects of* isc1*Δ cells can be partially rescued by* PGC1 *deletion*

Double deletion mutant *pgc1*Δ*isc1*Δ has been constructed as described in Materials and Methods. First we analyzed the mitochondrial phospholipid content in the newly constructed *pgc1*Δ*isc1*Δ strain to test whether it accumulated PG as expected. To do this, we compared phospholipid composition of mitochondrial membranes in the wild type, *isc1*Δ, *pgc1*Δ and *pgc1*Δ*isc1*Δ strains. As mentioned above, the transition of Isc1 from ER to mitochondria is controlled by Sch9 (10). To distinguish the effect of the complete absence of Isc1 from the effect of the absent change in its local distribution, we also included the *sch9*Δ mutant into this comparison.

Results of phospholipid content analysis are shown in Fig. 1. As expected, we detected a remarkable accumulation of PG in *pgc1*Δ*isc1*Δ cells. In fact, this mutant accumulated about twice as much PG as the *pgc1*Δ strain did. We did not identify any significant difference in cardiolipin content between the analyzed strains. However, we found significantly decreased PE content in mitochondria of *isc1*Δ strain. This drop in PE levels was not observed in *sch9*Δ cells, leading us to conclude that this was not due to loss of Isc1 translocation into mitochondria, as neither of these mutants contained mitochondrial fraction of the protein, but rather to a general lack of Isc1 protein function in *isc1*Δ cells. In addition, wild type-like PE levels could be restored by *PGC1* deletion in *isc1*Δ strain. In accordance with the previously published data, we also found lower PC content in the two strains lacking the *PGC1* allele (23), and a small but statistically significant decrease of PC levels was detected in *sch9*Δ strain, too.

**Figure 1.**
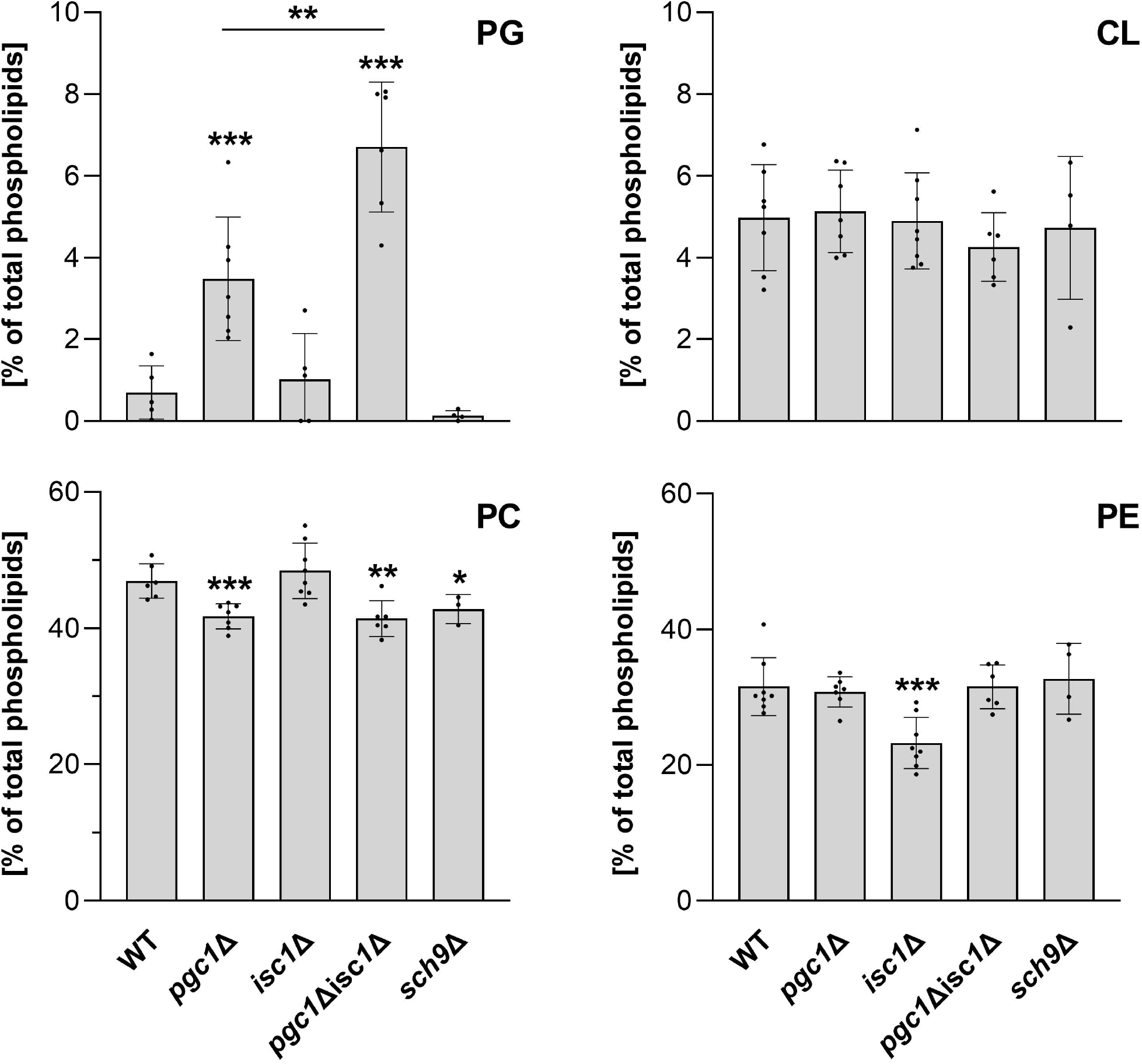
Phospholipid content analysis. Wild type, *isc1*Δ, *pgc1*Δ, *pgc1*Δ*isc1*Δ and *sch9*Δ strains of *S. cerevisiae* (see Supplementary Table for details) were cultivated in SMDGE medium for 24 h. Lipids from the mitochondrial fractions were extracted, separated, and the relative amounts of phospholipids were calculated based on the contents of inorganic phosphate. Data represent mean values from at least 4 independent experiments (dots) ± SD (errorbars). Statistically significant differences between mutant strains and wild type or between *pgc1*Δ*isc1*Δ and *isc1*Δ strain are indicated (asterisks; * – p < 0.05; ** – p < 0.01; *** – p < 0.001). CL, cardiolipin; PC, phosphatidylcholine; PE, phosphatidylethanolamine; PG, phosphatidylglycerol; WT, wild type.

Next we compared the mitochondrial function in the analyzed strains. Specifically, we measured the respiratory capacity in mitochondria isolated from the wild type, *isc1*Δ, *pgc1*Δ and *pgc1*Δ*isc1*Δ cells in the ADP-activated state in the presence of NADH (OXPHOS capacity). Consistent with previously published data, we detected OXPHOS capacity reduced to ∼20% of the wild type value in *isc1*Δ (17, 19, 24) and significantly increased in *pgc1*Δ strain (22, 23). In the *pgc1*Δ*isc1*Δ double mutant, OXPHOS capacity was partially restored to 76 ± 11% of wild type value (Fig. 2A). Cytochrome *c* reductase and cytochrome *c* oxidase (Complex III and IV, respectively) activities measured *in vitro* revealed that the observed changes in OXPHOS capacity are mainly due to changes in Complex IV activity. While no significant variations in Complex III activity were detected among the strains analyzed, Complex IV activity decreased to 18 ± 7% in *isc1*Δ mitochondria and was nearly doubled in *pgc1*Δ mitochondria. In the double mutant, Complex IV activity was restored to the level of the wild type strain (Fig. 2B, C). These differences in Complex IV activity only partially correlated with the observed differences in the amount of the Complex IV subunit, Cox4, between the strains analyzed. In particular, the recovery of Complex IV activity in the *pgc1*Δ*isc1*Δ double mutant cannot be explained by differences in protein abundance alone (Fig. 2D).

**Figure 2.**
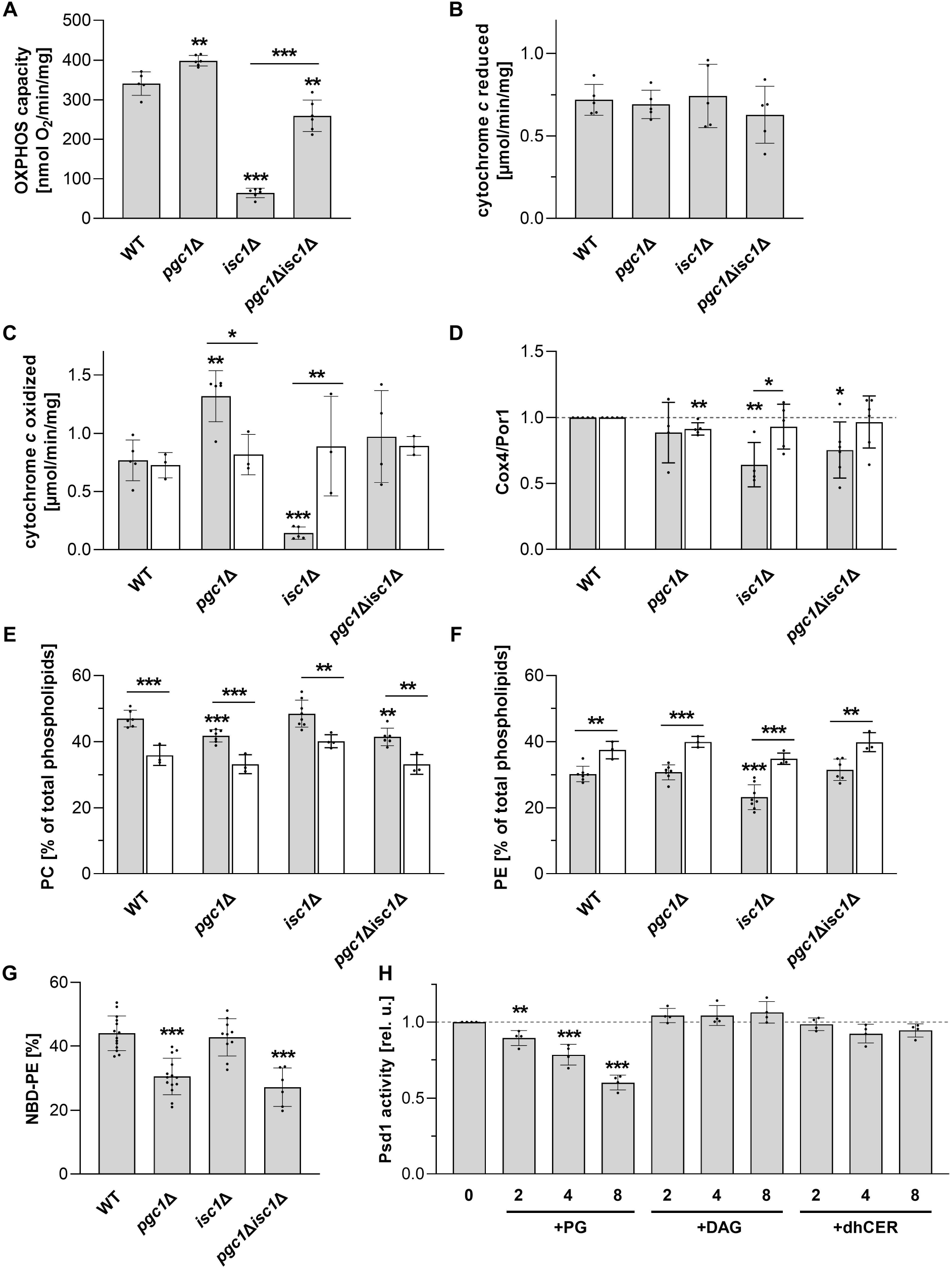
Role of Pgc1 and Psd1 in reduced respiration of *isc1*Δ cells. Wild type, *isc1*Δ, *pgc1*Δ and *pgc1*Δ*isc1*Δ strains of *S. cerevisiae* (see Supplementary Table for details) were cultivated in SMDGE medium for 24 h without (grey bars, at least 4 independent experiments) or with (white bars, 3 independent experiments) added ethanolamine (final concentration: 2 mM). Isolated mitochondria from these cells were used to measure oxygen consumption in the presence of ADP (OXPHOS capacity), with NADH as a respiratory substrate (A). *In vitro* activity of Complex III (B) and Complex IV (C) was measured in lysates prepared from isolated mitochondria (see Materials and Methods for details). Relative levels of Cox4 (subunit of Complex IV) were normalized to Por1 level (D). Lipids from the mitochondrial fractions were extracted, separated, and the relative amounts of PC (E) and PE (F) were calculated based on the contents of inorganic phosphate. *In vitro* phosphatidylserine decarboxylase activity of Psd1 was measured in mitochondrial fractions using NBD-PS as a substrate. Fluorescent product (NBD-PE) was quantified (G). The dependence of Psd1 activity on the concentration of PG, DAG, or dhCER (see Materials and Methods for details) in the reaction (numbers denote the amounts of added lipids in µg per reaction) was measured in mitochondrial fractions of the wild type cells (H). Data represent mean values from at least 4 independend experiments (dots) ± SD (errorbars). Statistically significant differences between mutant strains and wild type or between samples with and without added ethanolamine are indicated in A-G, statistically significant differences to the sample with no added lipids are indicated in H (asterisks; * – p < 0.05; ** – p < 0.01; *** – p < 0.001). WT, wild type; PG, phosphatidylglycerol; DAG, diacylglycerol; dhCER, dihydroceramide.

PE is a non-bilayer lipid which is, besides CL, important for mitochondrial structure and function (25, 26). The defect in *de novo* PE synthesis can be bypassed by the supply of exogenous ethanolamine during cultivation (27). We therefore tested whether ethanolamine supply can affect the *isc1*Δ phenotype. Comparison of the phospholipid profiles showed that for all strains analyzed, cells grown in medium supplemented with ethanolamine contained a higher PE fraction at the expense of the PC fraction (Fig. 2E, F). In addition, Complex IV activity in *isc1*Δ cells fed with ethanolamine was restored to wild type levels (Fig. 2C). We conclude that the deficiency of Complex IV in *isc1*Δ cells is due to a reduced PE fraction in their mitochondrial membranes. Similar observation that a moderate reduction in PE levels of <30% induces severe alterations in mitochondrial morphology and function of Complex IV has been reported previously in mice (28).

Both defects found in *isc1*Δ mutant, i.e., decreased PE content and drop in Complex IV activity, are also generated in cells lacking mitochondrial phosphatidylserine (PS) decarboxylase Psd1 (29), mediating the crucial step of the major pathway of PE synthesis (Schuiki and Daum, 2009). To determine whether the phenotype of *isc1*Δ cells is due to impaired Psd1 function, we compared the mitochondrial PS decarboxylase activity in the analyzed strains (Fig. 2G). No significant decrease of Psd1 activity could be detected in the mitochondrial fraction isolated from *isc1*Δ cells, which indicated that PS decarboxylase deficiency is not the primary reason for the *isc1*Δ phenotype. In contrast, we found significantly decreased Psd1 activity in mitochondrial fractions of both strains lacking *PGC1* (Fig. 2G), which did not affect the PE content in these cells (Fig. 1), and either normal or even increased activity of Complex IV was detected in mitochondrial fractions of *pgc1*Δ*isc1*Δ and *pgc1*Δ cells, respectively (Fig. 2C). Modulation of Psd1 function thus does not appear to be the cause of the rescue of the *isc1*Δ phenotype by *PGC1* deletion (Figs. 1, 2A-C).

We searched for the reason why the strains lacking *PGC1* gene exhibited decreased PS decarboxylase activity. *PGC1* deletion leads to massive PG accumulation (22, 31; Fig. 1). We therefore asked whether increased PG content could be sufficient to reduce the rate of PS decarboxylation. We measured *in vitro* PS decarboxylase activity in isolated wild type mitochondria (Psd1 activity) supplemented with different amounts of PG. As documented in Fig. 2H, Psd1 activity gradually decreased with increasing amounts of PG in reaction. We conclude that reduced PS decarboxylase activity documented in Fig. 2G was a consequence of PG accumulation in Pgc1 deficient cells.

### Diacylglycerol inhibits PG synthesis

Besides PS decarboxylase Psd1, another enzyme whose activity can be affected by the absence of Pgc1 is PGP synthase Pgs1. In an *in vitro* assay, we detected no difference in Pgs1 activity between the wild type and *pgc1*Δ strain. However, significantly reduced Pgs1 activity in *isc1*Δ cells was restored to the wild type levels by *PGC1* deletion (Fig. 3A). Of note, reduction in Pgs1 activity caused by the absence of *ISC1* was independent of the presence of Sch9, as Pgs1 activity comparable to the wild type level was detected in *sch9*Δ strain. Similarly, the restoration of Pgs1 activity following *PGC1* deletion occurred both in *isc1*Δ and *isc1*Δ*sch9*Δ background, i.e., again independently of Sch9. This suggests that Pgs1 activity in *isc1*Δ cells is reduced due to loss of Isc1 function in general, not due to a change in Isc1 cellular localization, which is thought to be driven by Sch9 (10).

**Figure 3.**
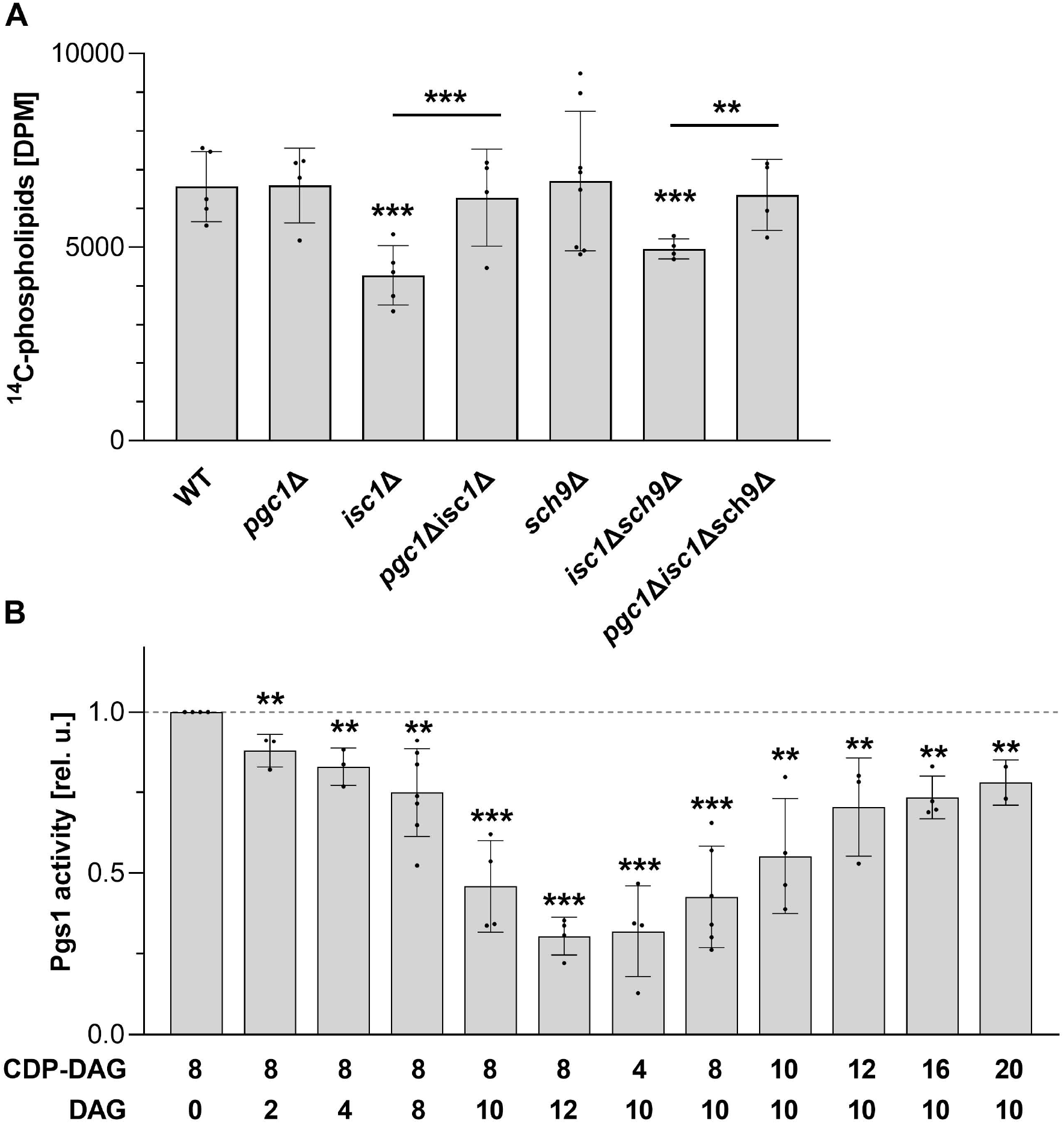
Regulation of Pgs1 activity. Indicated strains of *S. cerevisiae* (see Supplementary Table for details) were cultivated in SMDGE medium for 24 h. *In vitro* Pgs1 activity in mitochondrial fractions was measured by the amount of radioactive carbon ^14^C incorporated into phospholipids, using ^14^C-labeled glycerol-3-phosphate as a substrate (A; see Materials and Methods for details). The dependence of Pgs1 activity on the concentration of CDP-DAG, DAG or their combinations in the reaction was measured in mitochondrial fractions of the wild type cells (B; numbers denote the amounts of added lipids in µg per reaction). Data in A and B represent mean values from at least 3 independent experiments (dots) ± SD (errorbars). Statistically significant differences between mutant strains and the wild type, *pgc1*Δ*isc1*Δ and *isc1*Δ or between *pgc1*Δ*isc1*Δ*sch9*Δ and *isc1*Δ*sch9*Δ are indicated (asterisks; * – p < 0.05; ** – p < 0.01; *** – p < 0.001). WT, wild type. WT, wild type; DAG, diacylglycerol; CDP-DAG, cytidine diphosphate diacylglycerol.

The mechanism of the proposed ability of Pgc1 to modulate Pgs1 activity was investigated in more detail. Molecular function of Pgc1 is to hydrolyze PG to diacylglycerol (DAG) and glycerol-3-phosphate (31). Pgs1 transfers the phosphatidyl group from CDP-DAG to the hydroxyl group of glycerol-3-phosphate (32) to synthesize PGP, which is rapidly dephosphorylated to PG. In our *in vitro* analysis, we detected maximal Pgs1 activity in the presence of 8 µg CDP-DAG substrate (Fig. 3B). This activity gradually decreased upon addition of increasing concentration of DAG into the reaction. This negative correlation occurred specifically for DAG and did not occur upon addition of other lipids or upon increasing the amount of the substrate (CDP-DAG) alone. No lipid other than the substrate contributed to the Pgs1 activity. Negligible Pgs1 activity was detected in the absence of CDP-DAG (Supplementary Fig. S1A). Finally, the DAG-inhibited Pgs1 activity was recovered by addition of higher amounts of CDP-DAG, indicating the competition between the two compounds (Fig. 3B).

In this context, it is worth mentioning that recovery of Pgs1 activity by the *PGC1* deletion is not a new phenomenon. We reported recently that reduced Pgs1 activity in a strain lacking monolyso-CL transacylase Taz1 was fully recovered in a *pgc1*Δ*taz1*Δ double deletion mutant. Analogous to *pgc1*Δ*isc1*Δ, the *pgc1*Δ*taz1*Δ strain accumulated PG. Consistent with our recent data, it could be the lower DAG production associated with the *PGC1* deletion that led to the observed restoration of Pgs1 activity in *pgc1*Δ*taz1*Δ cells (23).

Taken together, our data suggest that the Pgc1-mediated PG hydrolysis product, DAG, is enough to reduce Pgs1 activity by competitively inhibiting CDP-DAG binding to the enzyme. DAG freely diffuses across biological membranes (33, 34) and could thus represent an effective tool for a cell to mediate both the signalization of increased PG degradation, i.e., reduced PG demand, and self-regulation of PG production by Pgs1.

### Functional interplay of phospholipases Pgc1 and Isc1

It has been documented that Isc1 is activated *in vitro* by the anionic phospholipids PG, CL and PS (35). We asked whether excess PG accumulated in cells lacking *PGC1* gene is sufficient to increase Isc1 activity in the *pgc1*Δ strain. Measurement of Isc1 activity *in vitro* showed that under conditions of respiratory growth, the opposite is true and Isc1 activity is significantly decreased in the whole cell homogenate as well as in the ER and mitochondrial membrane fractions isolated from the *pgc1*Δ cells, if compared to the wild type (Fig. 4A). To verify that the reduction in Isc1 activity observed in the *pgc1*Δ strain was indeed due to an excess of PG, we compared Isc1 activity in mitochondria isolated from these cells grown in medium with or without inositol. Inositol is a known repressor of PG synthesis (36, 37) and the cells with the *PGC1* gene deletion grown in inositol-supplemented medium contain similar amounts of PG as wild type (31). Consistent with this, we detected wild type Isc1 activity in mitochondria isolated from *pgc1*Δ cells grown in inositol-containing medium. Of note, the effect of PG-suppressed Isc1 activity was independent on the presence of Sch9 (Supplementary Fig. S1B).

**Figure 4.**
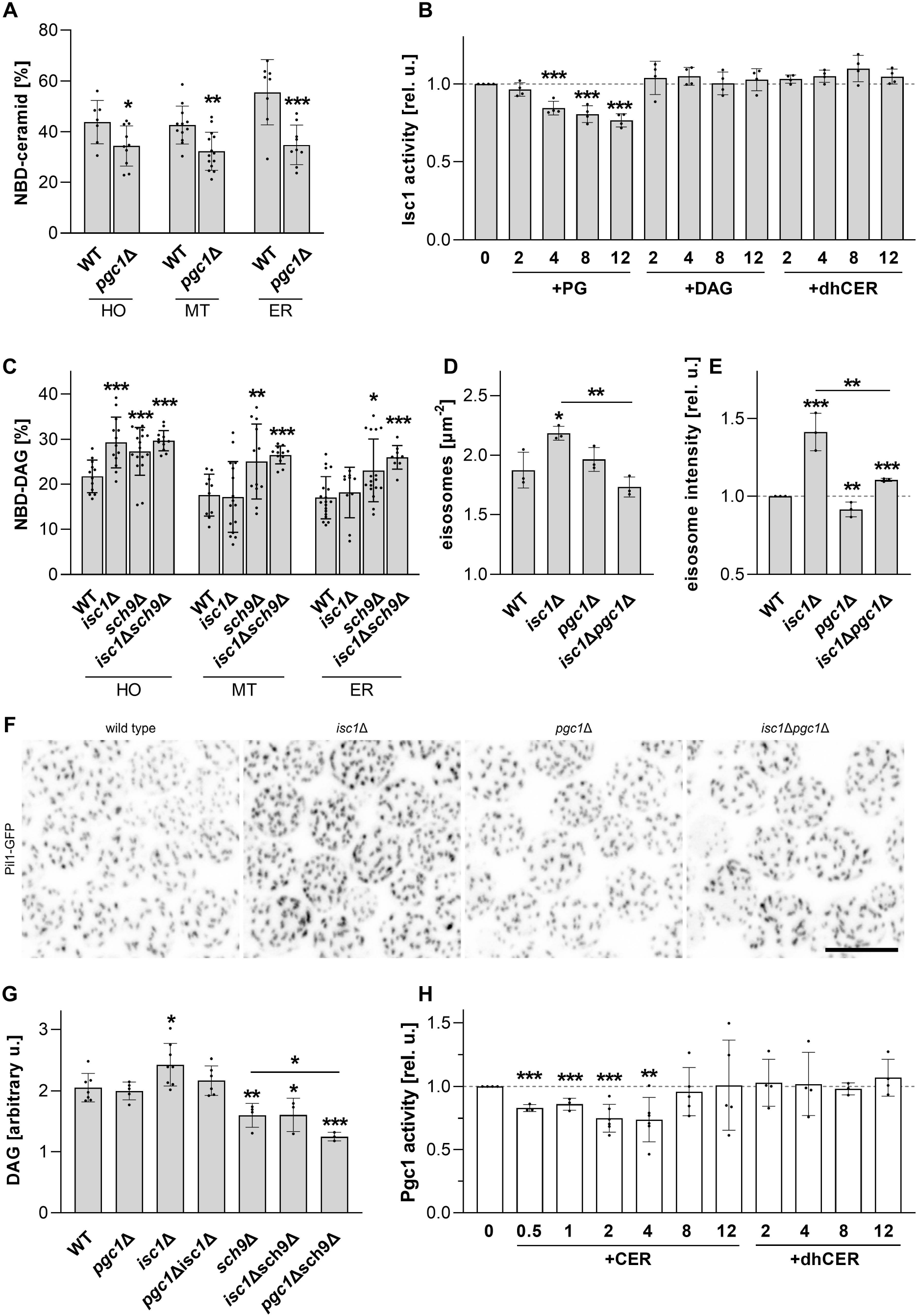
Regulation of Isc1 and Pgc1 activity. Indicated strains of *S. cerevisiae* (see Supplementary Table for details) were cultivated in SMDGE medium for 24 h. *In vitro* activities of Isc1 (A, B) and Pgc1 (C, H) were measured in whole cell homogenate (HO), isolated mitochondria (MT) and the ER fraction (ER; in A, C), or after addition of indicated amounts (in µg per reaction) of selected lipids into the mitochondrial fractions (B, H). In wild type, *isc1*Δ, *pgc1*Δ and *pgc1*Δ*isc1*Δ strains expressing Pil1-GFP (see Materials and Methods for details), structure of the plasma membrane-associated eisosomes was compared. Eisosome density and fluorescence intensity were analyzed (D, E) in maximum projections of six consecutive confocal sections (F). Bar: 5 μm. At least 250 cells/strain were analyzed in each experiment. Lipids from the mitochondrial fractions were extracted, separated, and the amount of DAG was compared (G). Data represent mean values from at least 4 independent experiments (dots) ± SD (errorbars). Statistically significant differences between mutant strains and wild type or between *pgc1*Δ*isc1*Δ and *isc1*Δ strain are indicated in A, C-E and G, statistically significant differences to the sample with no added lipid are indicated in B, H (asterisks; * – p < 0.05; ** – p < 0.01; *** – p < 0.001). WT, wild type; PG, phosphatidylglycerol; DAG, diacylglycerol; dhCER, dihydroceramide; CER, ceramide.

Similar effect of reduced Isc1 activity could be observed when excess PG was added into the membrane samples isolated from wild type cells. No change in Isc1 activity was observed upon addition of other lipids (Fig. 4B). The relative decrease of the Isc1 activity observed at higher concentrations of PG was comparable to the difference between the enzyme activities in mitochondria isolated from wild type and *pgc1*Δ cells (compare Fig. 4A, B). We conclude that Pgc1 participates in Isc1 activation by preventing PG accumulation. Similarly, the increase in Isc1 activity observed during the diauxic shift (19) can be interpreted as a release from PG inhibition under conditions of increased need for PG as a major precursor in CL biosynthesis at the onset of respiration.

Activity of Pgc1 itself is tightly regulated, too. Large pool of inactive Pgc1 is stored in lipid droplets and is activated only after release to the inner cellular membranes (ER, outer mitochondrial membrane; Kubalová *et al*, 2019), where it is also rapidly degraded by endoplasmic-reticulum-associated protein degradation (ERAD) pathway (38). The molecular mechanism of timely release of Pgc1 from the storage to the places of its enzymatic activity remains to be identified. However, since this step involves translocation of the enzyme from the lipid monolayer on the surface of the lipid droplet to the lipid bilayers of various subcellular membranes (20), it is reasonable to expect that it is directly influenced by the lipid composition of the target membranes. For example, Isc1-catalyzed hydrolysis of complex sphingolipids reduces these tightly exoplasmically localized lipids to ceramide, which diffuses between membrane leaflets (39). Thus, Isc1 activity can substantially alter the properties of the ER and/or outer mitochondrial membranes, in which active Pgc1 has been localized (20). We asked therefore whether Isc1 could play a role in regulation of Pgc1 function.

First we compared Pgc1 activity in the wild type and *isc1*Δ strains. We detected increased Pgc1 activity in the whole cell homogenate of the *isc1*Δ cells. This increase was not detected either in mitochondrial or ER fraction (Fig. 4C), which suggested that it could reflect a changed Pgc1 activity in other cellular compartments of *isc1*Δ mutant. In the genome-wide screen, *isc1*Δ cells exhibited hyper-assembled eisosomes at their plasma membrane (11). We checked whether this particular phenotype of the *isc1*Δ mutant could also be rescued by deletion of *PGC1*. To this end, we compared the eisosomal patterns in wild type, *isc1*Δ, *pgc1*Δ, and *pgc1*Δ*isc1*Δ cells, which were visualized using the fluorescently labeled eisosome organizer Pil1. Quantitative analysis revealed a statistically significant increase in eisosome density and Pil1-GFP fluorescence intensity in the *isc1*Δ mutant. For both parameters, this increase could be compensated by deletion of *PGC1* (Fig. 4D-F).

Consistent with the increased Pgc1 activity, we also detected increased levels of the Pgc1-catalyzed hydrolysis product, DAG, in *isc1*Δ cells. DAG levels returned back to the wild type value in *pgc1*Δ*isc1*Δ strain, indicating that Pgc1 activity was indeed responsible for the effect (Fig. 4G). Moreover, the increase in DAG correlated markedly with the decreased Pgs1 activity in the *isc1*Δ strain (Fig. 3A). We consider this as further evidence that physiological lipid levels (ceramides, DAG) are able to regulate the activity of the enzymes involved (Pgc1 and Pgs1, respectively). Contrary to our expectation, there was no statistically significant decrease in DAG content in the *pgc1*Δ strain compared to the wild type value. However, the situation was different in the absence of Sch9 protein. In *sch9*Δ cells, sphingolipid profiling showed reduced levels of many ceramide species (10). It was therefore not surprising that, similar to the *isc1*Δ cells lacking the ceramide-generating lipase, we detected increased Pgc1 activity in the *sch9*Δ strain. In contrast to *isc1*Δ, this increase was statistically significant in all cell fractions analyzed (Fig. 4C). The total DAG content in *sch9*Δ mitochondria was lower compared to the wild type and it could not be elevated in this strain even by deletion of the *ISC1* gene. However, it decreased further in the absence of Pgc1, suggesting that, in contrast to the wild type and similar to the *isc1*Δ strain, a considerable fraction of DAG in *sch9*Δ cells is a result of Pgc1-mediated hydrolysis of PG. This difference between the wild type on one, and *isc1*Δ and *sch9*Δ backgrounds on the other side, illustrates the versatility and adaptability of lipid-mediated enzyme regulation. While ceramide production by Isc1 was critical for Pgc1 activity in the wild type, in the absence of the protein in *isc1*Δ or in the environment with deregulated sphingolipid biosynthesis in the *sch9*Δ strain, ceramide levels were out of the range significantly regulating Pgc1 activity and the contribution of Isc1 in the latter case was not essential. Conversely, the regulatory potential of Pgc1-generated DAG was dominant in both mutant strains, where it effectively reduced Pgs1 activity, and became ineffective in the wild type strain (Fig. 3A).

Next we asked whether Pgc1 activity in isolates from *isc1*Δ cells could be attenuated to wild type levels by adding exogenous ceramide to the *in vitro* assay. Indeed, a significant decrease in Pgc1 activity was detected when small amounts of ceramide were added to the reaction, with the lowest Pgc1 activity detected after the addition of 2 µg of ceramide (Fig. 4H). Further increases in the amount of ceramide in the reaction led to a gradual disappearance of its inhibitory effect. The increasing variance among biological replicates of the high ceramide samples suggested that the addition of high ceramide amounts affected the structure of the membrane isolates. One possible explanation could be lipid phase separation. This effect was highly specific, as no significant change in Pgc1 activity was observed after addition of C16 dihydroceramide (dhCER, Fig. 4H) or other lipid species (Supplementary Fig. S1C). The differential effect of distinct ceramide species on the phenotype of *isc1*Δ cells has been described previously (e.g. 19), but further experiments will be required to explain the molecular details of the observed phenomenon.

Two different type C phospholipases, Isc1 and Pgc1, share the subcellular localization at the ER and outer mitochondrial membrane (9, 20). In this study, we have shown that there is also a functional connection between them, because the activity of one influences the activity of the other and *vice versa*. In this feedback loop, the ceramide generated by Isc1 activity suppresses degradation (Fig. 4C) and stimulates the synthesis (Fig. 3A) of the CL precursor, PG, by inhibiting the enzymatic activity of Pgc1 and subsequently Pgs1. Feedback is provided by the PG itself, because higher levels of this particular lipid inhibit Isc1 activity (Fig. 5). This direct link between mitochondrial lipid metabolism and the sphingolipid biosynthesis pathway, which responds to a wide range of environmental cues (40, 41), could be of high physiological relevance.

**Figure 5.**
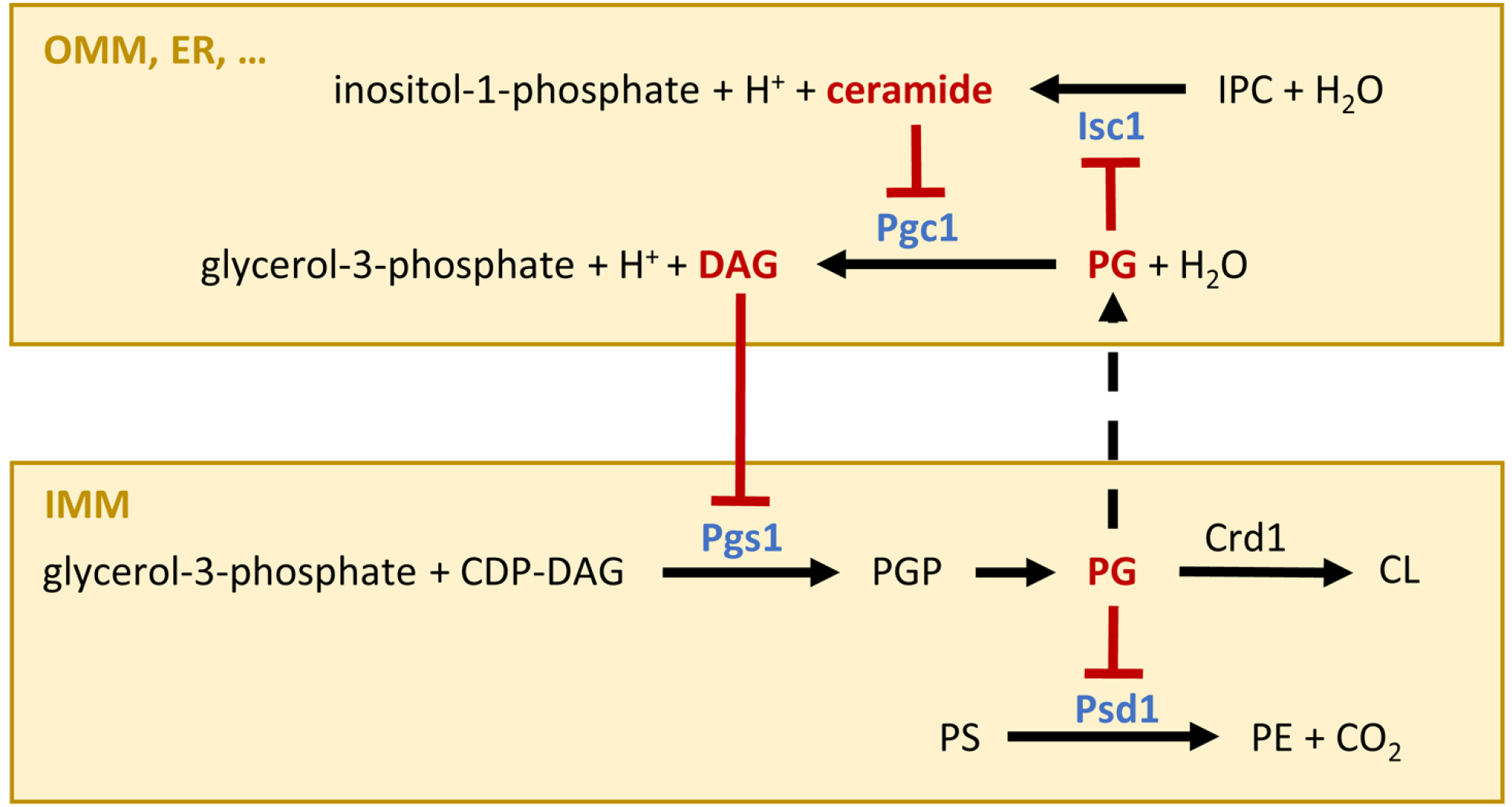
Mechanism of mutual regulation of Isc1 and Pgc1 phospholipases. Our data indicate that two lipids, DAG and PG, inhibit the key enzymes of mitochondrial phospholipid biosynthesis: (i) PGP synthase Pgs1, (ii) phosphatidylserine decarboxylase Psd1, and (iii) inositolphosphosphingolipid phospholipase C, Isc1. The latter is of particular importance because ceramide, a product of Isc1-catalyzed hydrolysis, inhibits the PG-specific phospholipase C, Pgc1, which degrades PG to form DAG. The entire loop ensures efficient control of PG (CL) and PE synthesis in the context of environmental stress stimuli affecting sphingolipid biosynthetic pathway. OMM – outer mitochondrial membrane, IMM – inner mitochondrial membrane.

In addition to the regulation of CL biosynthesis, actual PG levels may also affect the production of PE (Fig. 2G, H), another non-bilayer lipid that plays a key role in mitochondrial morphology and function and whose biological relevance largely overlaps with CL (26, 42). This type of PE regulation could be important in CL deficiency. For example, consistent with previously published data (43), we detected elevated PE levels in mitochondria of *crd1*Δ strain, which completely lacks CL (Supplementary Fig. S2). The absence of Crd1 leads to PG accumulation, but this pool of PG is not accessible to Pgc1-mediated hydrolysis (22). It has been suggested that in the absence of CL, PG fulfills its function in binding respiratory complexes (e.g. 44). Thus, it is likely that this bound PG cannot inhibit Psd1, which could account for the increase in PE levels. Unlike CL, cylindrical molecules of PG cannot provide the membrane curvature necessary for the formation of proper mitochondrial morphology. Elevated non-bilayer PE, molecules of which adopt an inverted cone shape due to a small ethanolamine headgroup, could complement this feature and together with PG participate in the functional replacement of the missing CL. It has been reported previously that further PG accumulation in the absence of CL, induced by *PGC1* deletion in the *crd1*Δ background, has aggravating effect on mitochondrial morphology (22). Within the suggested model (Fig. 5), the explanation is obvious: excess PG cannot bind to respiratory complexes for stoichiometric reasons. Instead, it becomes available not only for degradation by Pgc1 (22), but also for inhibition of Psd1. Consistent with this explanation, we did not observe increased levels of PE in the *pgc1*Δ*crd1*Δ strain (Supplementary Fig. S2). This lack of PE likely accounts for the increased frequency of aberrant mitochondrial sheets in this strain compared to *crd1*Δ (22).

The proposed model of lipid-mediated regulation of mitochondrial phospholipid homeostasis (Fig. 5) represents a simple way for the cell to ensure optimal mitochondrial structure and function in response to current needs. The fact that its functionality is based purely on the spontaneous movement of lipids along a concentration gradient, whether by lateral diffusion or intermembrane exchange, means, among other things, that the coupling of individual regulatory steps does not require energy and works under any conditions - nutrient deprivation, stress, etc. This makes it a universally applicable principle of cellular regulation.

## Materials and Methods

### Yeast strains and growth conditions

All *S. cerevisiae* strains used in this study are listed in Supplementary Table. Cell cultures were grown in complex media (YPD; 2 % peptone, 1 % yeast extract, 2 % glucose). For experiments, cells were grown aerobically at 30 °C for 24 h to diauxic shift in a defined synthetic medium prepared as previously described (SMDGE; 0.2 % glucose, 3% glycerol, and 1% ethanol as a carbon source; (45)). If not stated otherwise, SMDGE medium lacked inositol.

### Strains construction

Mutant strains *pgc1*Δ*::HIS3* of both mating types were prepared by replacement of *pgc1*Δ*::KanMX4* disruption cassette with *pgc1*Δ*::HIS3*, and mutant strain *sch9*Δ*::NatMX4* was prepared by replacement of *sch9*Δ*::KanMX4* disruption cassette with *sch9*Δ*::NatMX4* in the collection strains *pgc1*Δ, alpha-*pgc1*Δ, and *sch9*Δ, respectively, as described in (46). Double mutants *pgc1*Δ*isc1*Δ and *isc1*Δ*sch9*Δ were prepared by the transformation of *pgc1*Δ*::HIS3* and *sch9*Δ*::NatMX4*, respectively, with a disruption cassette *isc1*Δ*::KanMX4*, PCR amplified from *isc1*Δ strain. Yeast transformation was performed by the lithium acetate method (47). Triple mutant *pgc1*Δ*isc1*Δ*sch9*Δ was selected by tetrad analysis after crossing of two parental strains alpha-*pgc1*Δ*::HIS3* and *isc1*Δ*sch9*Δ as isogenic to BY4741 except for the three required deletions. To visualize Pil1-GFP *in vivo*, the wild type, *isc1*Δ, *pgc1*Δ, and *pgc1*Δ*isc1*Δ strains were transformed with the YIp128-*PIL1-GFP* plasmid. The plasmid was constructed by inserting the *PIL1* gene as a *Hind*III-*Bam*HI fragment into the YIp128-GFP integrative plasmid. Prior to transformation, the plasmid was linearized by cleavage with *Xba*I to enable its genome integration. Successful transformants were selected based on loss of leucine auxotrophy due to the *PIL1::GFP::LEU2* insert.

### Enzymatic assays

Intact mitochondria were isolated from the cells grown to diauxic shift as described previously (22). The final mitochondrial pellet was suspended in the respiration buffer (0.6 M mannitol, 20 mM HEPES/KOH pH 7.1, 2 mM MgCl2, 1 mM EGTA, 0.1 % fatty acid-free bovine serum albumin, 10 mM KH2PO4) and used for measurement of O2 consumption (OXPHOS capacity) as described in (23), the activity of cytochrome *c* reductase and the activity of cytochrome *c* oxidase was measured as described in (22).

Activity of Pgc1 was measured as described previously (20) with modifications in used amount of isolated subcellular fractions. Briefly, the mixture containing 40 μl of 0.3 M Tris/HCl, pH 7.4; 20 μl of NBD-PG (0.8 μg) and an the respective subcellular fraction (cell homogenate, mitochondrial or ER fraction in the amount corresponding to 50 μg of proteins) was incubated at 30°C for 40 min. Linearity of the reaction under these conditions has been verified earlier (Kubalova et al., 2019). The reaction was stopped by the addition of chloroform/methanol/HCl (2 ml; 100:100:0.6 v/v/v) and 1 ml of water. After the separation, the organic phase was dried under a stream of nitrogen. NBD-labeled lipids were separated by a one-dimensional thin layer chromatography for 15 min using a chloroform/methanol developing solvent (70:35; v/v). Degradation of exogenous fluorescent substrate NBD-PG to NBD-DAG was quantified using TLC scanner (Camag; 460 nm), and the phospholipase C activity of Pgc1 was determined as a fraction of NBD-DAG in the total NBD-labeled lipids.

Activity of Pgs1 was determined by quantification of the incorporation of radiolabeled substrate [^14^C]glycerol-3-phosphate into chloroform-soluble products as described previously (23). Briefly, the reaction mixture containing 50 mM MES-HCl pH 7.0, 0.1 mM MnCl2, synthetic lipids in indicated amounts (if not stated otherwise, 0.083 mM CDP-DAG has been used which corresponded to 8 µg of the lipid per reaction), 1 mM Triton X-100, 0.02 mM [^14^C]glycerol-3-phosphate (40,000 CPM/nmol), mitochondrial fraction corresponding to 25 μg of mitochondrial protein in a total volume of 120 μl was incubated for 20 min at 30 °C. The reaction was stopped by the addition of excess chloroform:methanol:HCl mixture (100:100:0.6), phase separation was then induced by water. Aliquots of the organic layer were evaporated under the stream of nitrogen, dissolved in scintillation mixture and the radioactivity of each sample was determined with a scintillation counter.

Isc1 activity was determined by quantification of the fluorescently labeled product, NBD-ceramide, after incubation of different cell fractions with fluorescently labeled substrate, NBD-sphingomyelin. Briefly, fresh cell fractions (cell homogenate 20 μg of proteins, mitochondrial fraction 15 μg of proteins and microsomal fraction enriched by ER 10 μg of proteins) were incubated in 100 μl of buffer containing 100 mM Tris/HCl pH 7.4, 5 mM MgCl2 and 0.5 μg of NBD-sphingomyelin. The mixture was incubated at 30 °C for 20 min. The reaction was stopped by the addition of chloroform/methanol/HCl (2 ml; 100:100:0.6; v/v/v) and 1 ml of water. After the separation, the organic phase was dried under a stream of nitrogen. NBD-labeled lipids were separated by a one-dimensional thin layer chromatography for 15 min using a chloroform/methanol developing solvent (70:35; v/v). The separated NBD-lipids were scanned with a TLC scanner (Camag) in the fluorescence mode at wavelength 460 nm, and the phospholipase activity of Isc1 was determined as a fraction of NBD-ceramide, a product of NBD-sphingomyelin hydrolysis, in the total NBD-lipids.

Activity of Psd1 was determined by quantification of fluorescently labeled product NBD-PE after incubation of the mitochondrial fraction with NBD-PS as substrate. The 140 μl of reaction mix contained 100 mM Tris/HCl pH 7.4, 10 mM EDTA, 0.8 μg NBD-PS and cell homogenate (40 μg of proteins) or mitochondrial fraction (15 μg of proteins). After the 20 min incubation at 30 °C the reaction was stopped, separated and quantified in same way as was described previously for Isc1 activity. Overall decarboxylase activity was determined as a fraction of NBD-PE in the total NBD-lipids.

### Synthetic lipids

In the *in vitro* assays testing the dependence of activity of various enzymes on specific lipids, the following synthetic lipid molecules (Merck) were added to the reaction: 1-palmitoyl-2-oleoyl-*sn*-glycerol (DAG; Cat. #800815), N-octanoyl-D-*erythro*-sphingosine (CER; Cat. #860645), N-palmitoyl-D-*erythro*-sphinganine (dhCER, Cat. #860634), 1-palmitoyl-2-oleoyl-*sn*-glycero-3-phospho-(1′-*rac*-glycerol) (PG; Cat. #810218), 1,3-dihexadecanoyl-2-(*cis*-9-octadecenoyl)glycerol (TAG, Cat. # D2157). Lipids were solubilized in chloroform:methanol (2:1, v:v). Before use, the required amount of lipids in the solution was added to the pyrex tube and dried under a stream of nitrogen. The dried lipids were resuspended in the appropriate buffer described above.

### Fluorescence microscopy

Yeast cells in a culture grown in SMDGE medium for 24 h at 30 °C were concentrated by brief centrifugation, immobilized on a 0.17 mm cover glass by a thin film of 1% agarose prepared in 50 mM potassium phosphate buffer (pH 6.3) and observed using LSM 880 (Zeiss) laser scanning confocal microscope equipped with a 100x PlanApochromat oil-immersion objective (NA = 1.4). The fluorescence signal of GFP (excited by 488 nm line of Ar laser) was detected using a photomultiplier tube after filtering with a bandpass 493–550 nm emission filter. 3D stacks of confocal sections were acquired with sampling in axial direction of 0.7 µm. Maximum intensity projections were calculated and quantitative analysis of confocal images was performed in ImageJ 1.53c software (ImageJ, U. S. National Institutes of Health, Maryland, USA) using custom-made macros developed for this study (available at github.com/jakubzahumensky/Isc1_paper; the repository also contains sample images).

### DAG analysis

Lipids from mitochondrial fraction prepared by zymolyase treatment (corresponding to 1 mg of proteins) were extracted, dried under nitrogen stream, resuspended in chloroform:methanol mixture (2:1), and separated by thin-layer chromatography (TLC) as described (48). Following the sulphuric acid stain, relative lipid content was determined using CAMAG WinCATS software after scanning TLC plates on CAMAG TLC scanner 3 at 475 nm.

### Miscellaneous

Preparation of microsomal fraction enriched by ER and phospholipid analysis was performed as described previously (20). Western blot analysis of Cox4 abundance was performed as described previously (22). Briefly, isolated mitochondria were lysed and separated on 12% denaturing polyacrylamide gel and blotted onto a nitrocellulose membrane. The membrane was blocked with 5% milk in TBS buffer (50 mM Tris/HCl pH 8.0, 150 mM NaCl, 0.05% (v/v) Tween 20) overnight and immunostained using primary rabbit anti-Cox4 (1:1000) or anti-Por1 (dilution 1:5000) for mitochondrial porin and secondary anti-rabbit (Sigma) antibodies. Secondary antibody was visualized using ECL+ kit (Amersham). Statistical comparisons were carried out by one-way analysis of variance using SigmaPlot 14 software (Systat Software, San Jose, CA). All graphs (GraphPad Prism 9 software, GraphPad Software, San Diego, CA) show the mean ± SD.

## Acknowledgments

The work was supported by the Scientific Grant Agency of the Ministry of Education, Science, Research and Sport of the Slovak Republic and the Slovak Academy of Sciences [grant number 2/0030/22] to MB, the Slovak Research and Development Agency contract [grant number APVV-20-0129] to MB, and the Czech Science Foundation projects [grant numbers 19–04052S, 20-04987S] to JM, PV and JZ.

## Author Contributions

Conceptualization, Funding acquisition, Methodology, Resources, Supervision, Writing – review & editing – MB, JM; Formal Analysis, Investigation; Validation – MB, LB, ID, PV, PK, JZ, JM; Software – JZ; Visualization – PV; Writing – original draft – JM

## Supplementary Material

**Figure S1 Further indications for regulation of Pgs1, Isc1 and Pgc1 activity**. (A) Dependence of Pgs1 activity on the concentration of CDP-DAG, DAG, TAG, PG, dhCER, or CER (numbers denote the amounts of added lipids in µg per reaction, see Materials and Methods for details) in the reaction was measured in mitochondrial fractions isolated from the wild type cells cultivated in SMDGE medium for 24 h. Data represent mean values from at least 2 independent experiments (dots) ± SD (errorbars). Relative values normalized to the sample with the addition of 8 µg CDP-DAG are presented. (B) Regulation of Isc1 activity by PG *in vivo*. Wild type, *pgc1*Δ, *sch9*Δ and *pgc1*Δ*sch9*Δ strains of *S. cerevisiae* (see Supplementary Table for details) were cultivated in SMDGE medium without (SMDGE I-; grey columns) or with the addition of 75 μM inositol (SMDGE I+; white) for 24 h. *In vitro* activities of Isc1 were measured in isolated mitochondria. Data represent mean values from at least 6 independent experiments (dots) ± SD (errorbars). Statistically significant differences between the mutants and the wild type and/or between the cells grown without or with inositol are indicated (asterisks; * – p < 0.05; ** – p < 0.01; *** – p < 0.001). WT, wild type. (C) Dependence of Pgc1 activity on the concentration of PG or DAG (numbers denote the amounts of added lipids in µg per reaction, see Materials and Methods for details) in the reaction was measured in homogenate of the *isc1*Δ cells cultivated in SMDGE medium for 24 h. Data represent mean values from at least 4 independent experiments (dots) ± SD (errorbars).

**Figure S2 Elevated PE content in single deletion mutant lacking CL synthase Crd1**. Wild type, *pgc1*Δ (see Supplementary Table for details), *crd1*Δ (*MAT*a *his3 leu2 met15 ura3 crd1::KanMX*) and *pgc1*Δ*crd1*Δ (*MAT*a *leu2 ura3 met15 crd1::KanMX pgc1::HIS3*) strains of *S. cerevisiae* were cultivated in SMD (synthetic medium as described in Materials and Methods, with 2% glucose as a sole carbon source) without inositol for 24 h. Lipids from the mitochondrial fractions were extracted, separated, and the relative amounts of phospholipids were calculated based on the contents of inorganic phosphate. Data represent mean values from at least 3 independent experiments (dots) ± SD (errorbars). Statistically significant differences between mutant strains and wild type or between *pgc1*Δ*crd1*Δ and *crd1*Δ strain are indicated (asterisks; * – p < 0.05; ** – p < 0.01; *** – p < 0.001). CL, cardiolipin; PA, phosphatidic acid; PC, phosphatidylcholine; PE, phosphatidylethanolamine; PG, phosphatidylglycerol; PI, phosphatidylinositol; PS, phosphatidylserine; WT, wild type.

**Supplementary Table Yeast strains used in this study**.

