## Supplementary figures and images for "Two different phospholipases C, Isc1 and Pgc1, cooperate to regulate mitochondrial function"

### Supplemental Figure S1

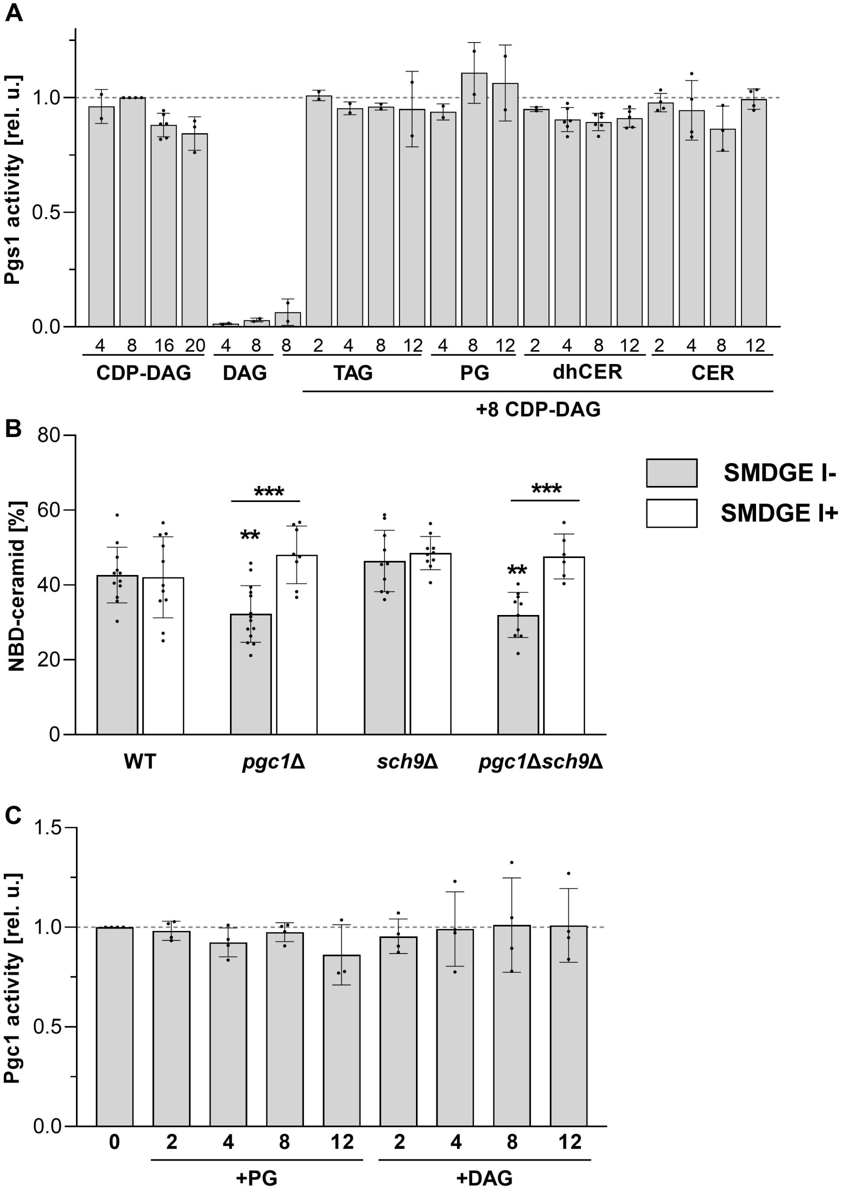

### Supplemental Figure S2

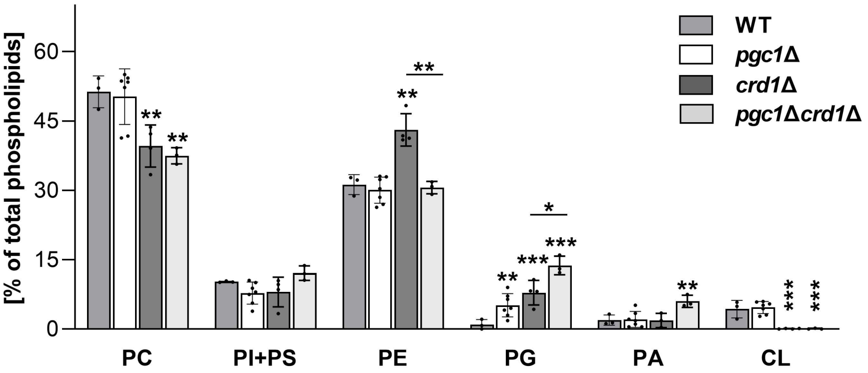
