## Supplemental Table for "Two different phospholipases C, Isc1 and Pgc1, cooperate to regulate mitochondrial function"

**Supplementary Table. Yeast strains used in this study.**

| Strain | Genotype | Source |
|---|---|---|
| BY4741, wild type (WT) | *MAT*a *his3Δ1 leu2Δ0 met15Δ0 ura3Δ0* | Euroscarf |
| BY4742 | *MATα his3Δ1 leu2Δ0 lys2Δ0 ura3Δ0* | Euroscarf |
| *pgc1Δ* | BY4741; *pgc1Δ::KanMX4* | Euroscarf |
| *isc1Δ* | BY4741; *isc1Δ::KanMX4* | Euroscarf |
| *sch9Δ* | BY4741; *sch9Δ::KanMX4* | Euroscarf |
| alpha-*pgc1Δ* | BY4742; *pgc1Δ::KanMX4* | Euroscarf |
| *pgc1Δ::HIS3* | BY4741; *pgc1Δ::HIS3* | This study |
| alpha-*pgc1Δ::HIS3* | BY4742; *pgc1Δ::HIS3* | This study |
| *sch9Δ::NatMX4* | BY4741; *sch9Δ::NatMX4* | This study |
| *pgc1Δisc1Δ* | BY4741; *pgc1Δ::HIS3, isc1Δ::KanMX4* | This study |
| *isc1Δsch9Δ* | BY4741; *isc1Δ::KanMX4, sch9Δ::NatMX4* | This study |
| *pgc1Δsch9Δ* | BY4741; *pgc1Δ::HIS3, sch9Δ::KanMX4* | This study |
| *pgc1Δisc1Δsch9Δ* | *MAT*a *his3Δ1 leu2Δ0 met15Δ0 ura3Δ0*; *pgc1Δ::HIS3, isc1Δ::KanMX4, sch9Δ::NatMX4* | This study |
